# RAB-5 regulates regenerative axonal fusion by controlling EFF-1 endocytosis

**DOI:** 10.1101/339150

**Authors:** Casey Linton, Brent Neumann, Rosina Giordano-Santini, Massimo A. Hilliard

## Abstract

Following a transection injury to the axon, neurons from a number of species have the ability to undergo spontaneous repair via fusion of the two separated axonal fragments. In the nematode *C. elegans*, this highly efficient regenerative axonal fusion is mediated by Epithelial Fusion Failure-1 (EFF-1), a fusogenic protein that functions at the membrane to merge the two axonal fragments. Identifying modulators of axonal fusion and EFF-1 is the next step towards harnessing this process for clinical applications. Here, we present evidence that the small GTPase RAB-5 acts to inhibit axonal fusion, a function achieved via endocytosis of EFF-1 within the injured neuron. Consequently, we find that perturbing RAB-5 activity increases the capacity of the neuron to undergo axonal fusion, through enhanced membranous localization of EFF-1 and the production of extracellular EFF-1-containing vesicles. These findings identify RAB-5 as a novel regulator of axonal fusion and the first regulator of EFF-1 in neurons.

---

Treating nerve injuries is of great interest in the clinical setting. Significant research efforts have been dedicated to developing new methods to repair an axon following transection, as current outcomes are less than optimal. A novel approach to this problem has come from work in invertebrate systems, where some neurons are capable of spontaneously rejoining the two separated axonal fragments after transection. This regenerative mechanism, called axonal fusion, has been observed in a number of species^1–5^, but has been best characterized in the mechanosensory neurons of the nematode *C. elegans*^6–10^. To undergo axonal fusion, the proximal axon (still attached to the cell body) must regrow, reconnect, and then fuse with its separated axonal fragment. We and others have shown in *C. elegans* that this fusion not only restores continuity of the axon following UV laser axotomy^7^, but also restores neuronal function^9,10^.

The key molecular effector of axonal fusion is the *C. elegans* membrane fusogen EFF-1. EFF-1, a nematode-specific transmembrane glycoprotein with structural and functional similarity to class II viral fusion proteins^11^, functions to merge closely apposed plasma membranes. EFF-1 activity in the injured neuron is crucial for successful axonal fusion; *eff-1* mutant animals exhibit severe axonal fusion defects^6,8^, which can be rescued by expression of wild-type EFF-1 selectively in the injured neuron, revealing its cell-autonomous function^8^. There is strong evidence that the activity of EFF-1 is controlled via its dynamic subcellular localization. To mediate fusion, EFF-1 must be inserted into the membrane, and it is inactive when sequestered in intracellular compartments. This was first demonstrated in other cell types in *C. elegans*, including the hypodermis, where EFF-1 mediates cell-cell fusion for the formation of syncytia during development^12^. In these cells, EFF-1 is mobilized from intracellular compartments to the plasma membrane where it mediates fusion^13,14^. Similarly, in the Posterior Lateral Microtubule (PLM) mechanosensory neurons, EFF-1 exists largely within puncta in the steady state, but following injury is mobilized to the regenerating axonal membrane^8^. How these changes in EFF-1 localization and activity are regulated within the neuron is currently unknown, but they likely represent a critical step in the control of axonal fusion.

To date, the only other molecules implicated in *C. elegans* axonal fusion are components of the apoptotic clearance machinery^8^ and the conserved microRNA *let-7*, which functions to regulate this pathway^10^. The clearance signalling pathways were first described as a mechanism for apoptotic cell corpse engulfment^15–19^, but appear to have been repurposed for axonal fusion, and function upstream of EFF-1 to promote recognition of the severed axonal fragment. Axonal injury triggers exposure of the phospholipid phosphatidylserine (PS) on the surface of the severed axon. PS acts as a ‘save me’ signal by recruiting both the PS receptor PSR-1, present on the growth cone of the regrowing fragment, and the secreted PS-binding protein TTR-52, which initiate signalling to improve the efficiency of axonal fusion^8,9^. Animals with mutated *psr-1* or *ttr-52* exhibit axonal fusion defects, which can be rescued with overexpression of EFF-1, indicating that EFF-1 acts genetically downstream of these recognition molecules^8^.

However, axonal fusion must also involve additional, undiscovered molecules. In the absence of the apoptotic genes, axonal fusion can still occur at a low rate, and EFF-1 is still mobilized to the membrane, indicating the existence of mechanisms which can bypass the apoptotic recognition machinery to enable fusion. Here, we identify a novel regulator of axonal fusion. We reveal that the GTPase RAB-5 can negatively regulate axonal fusion functioning within the injured neuron, and present evidence that this protein controls the level of EFF-1 on the neuronal membrane via endocytosis. Thus, we propose a model in which recycling of EFF-1 via RAB-5 is a critical mechanism for the control of fusogen function and axonal fusion as a mechanism of repair.

## RESULTS

### RAB-5 regulates the rate of regenerative axonal fusion

To identify molecules with a role in axonal fusion, we used mutations in specific genes and tested their effect on the rate of axonal fusion in a *psr-1* mutant background. This sensitized background allowed us to assess whether a candidate gene could effectively bypass the apoptotic recognition machinery and modulate axonal fusion rates through an alternative pathway.

Our primary candidate was the endocytic GTPase RAB-5, as this protein has been shown to modulate the rates of fusion in other *C. elegans* tissues. Specifically, loss-of-function or depletion of RAB-5 leads to a hyperfusion phenotype in the *C. elegans* hypodermis^14^. We first investigated the effect of perturbing RAB-5 activity by performing axotomies on the mechanosensory neurons of *psr-1* mutant animals expressing a dominant negative version of RAB-5 (RAB-5(DN)). Axotomies were performed on the PLM neurons, approximately 50 μm anterior to the cell body, as previously described^7,8^. The rate of axonal fusion was calculated from axons that regrew and visibly reconnected to their distal fragment. Successful axonal fusion occurred if the distal fragment was maintained 48 h post-axotomy (Fig. 1a); fusion was considered unsuccessful if the distal fragment instead underwent degeneration (Fig. 1b).

**Figure 1.**
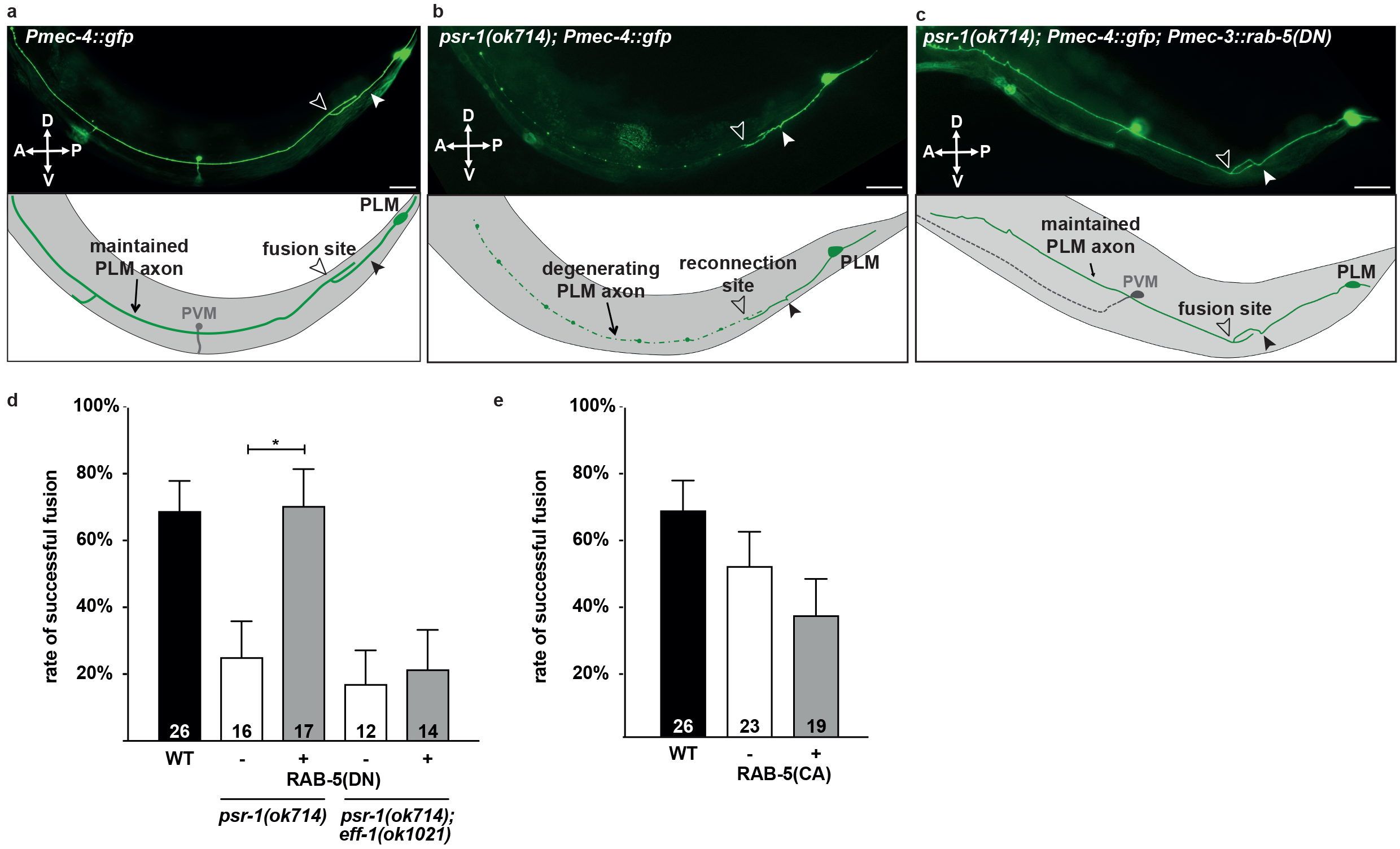
Perturbing RAB-5 activity increases EFF-1-dependent axonal fusion rates. **a** Successful PLM axonal fusion in a wild-type animal (*Pmec-4::GFP*) 48 h post-axotomy. The filled arrowhead indicates the cut site; the open arrowhead indicates the site of fusion. Another neuron (PVM, grey in schematic) is visible in the mid body of the animal. **b** Defective PLM axonal fusion in a *psr-1(ok714)* animal, 48 h post-axotomy. The filled arrowhead indicates the cut site; the open arrowhead indicates the site of reconnection (which did not result in fusion). **c** Rescue of PLM axonal fusion in a *psr-1(ok714)* animal by expression of RAB-5(DN), 48 h post-axotomy. The filled arrowhead indicates the cut site; the open arrowhead indicates the site of fusion. PVM (grey in schematic) can be seen but is out of focus. **d** Quantification of axonal fusion in *psr-1(ok714)* animals expressing RAB-5(DN) in the PLM neuron, demonstrating rescue of the *psr-1(ok714)* axonal fusion defect. This rescue is suppressed in *psr-1(ok714); eff-1(ok1021)* mutant animals, indicating that the effect of RAB-5(DN) is *eff-1*-dependent. **e** Quantification of axonal fusion in animals expressing RAB-5(CA) in the PLM neuron shows no significant effect on the level of axonal fusion compared to controls. Error bars indicate the standard error of proportions. n values are listed below each bar. P values from t-test: *P<0.05. D = dorsal, V = ventral, A = anterior, P = posterior for all images. Scale bars 25 μm.

As previously shown, *psr-1* mutant animals displayed a defective rate of axonal fusion compared with wild-type animals^8^. Remarkably, we found that expression of RAB-5(DN) in the PLM neuron was sufficient to rescue this fusion rate to wild-type levels (Fig. 1c, d). We also performed axotomies on wild-type animals expressing constitutively active RAB-5 (RAB-5(CA)) in the mechanosensory neurons, hypothesizing that this increased RAB-5 activity might generate the opposite phenotype and reduce the rate of axonal fusion. However, although there was a trend towards reduction, there was no significant change in the axonal fusion rates in these animals (Fig. 1e).

Given that EFF-1 is the key molecular effector of this process, we next asked whether RAB-5 modulates axonal fusion through an interaction with EFF-1. To address this, we performed axotomies on double mutant animals of *eff-1* and *psr-1* expressing RAB-5(DN). We found that loss of *eff-1* suppressed the increase in axonal fusion mediated by RAB-5(DN) (Fig. 1d), indicating that RAB-5(DN) increases the axonal fusion rate in an *eff-1*-dependent manner.

### RAB-5 controls EFF-1 localization to the plasma membrane

We next asked how RAB-5 was regulating the activity of EFF-1 in mediating axonal fusion. We suspected that perturbing RAB-5 prevented the endocytosis of EFF-1, which would increase the amount of EFF-1 available at the membrane and enhance its activity. RAB-5 and its mammalian orthologue Rab5 are known to localize to early endosomes and play important roles in endocytosis. They facilitate the transport of clathrin-coated vesicles to early endosomes, fusion between endosomes, and cargo trafficking from endosomes into lysosomes for degradation^20–24^. Correspondingly, altering RAB-5 activity produces specific cellular phenotypes: perturbing RAB-5 activity using RAB-5(DN) inhibits endocytosis and causes membranous accumulation of proteins^25^, whereas expression of constitutively active RAB-5(CA) leads to excessive early endosome fusion and the presence of enlarged early endosomes^25,26^. In the *C. elegans* hypodermis, depletion or loss-of-function of RAB-5 results in mis-localization of EFF-1 to the plasma membrane, which in turn is associated with excessive hypodermal cell-cell fusion^14^. We therefore predicted that perturbing RAB-5 activity in the PLM neurons would result in similar changes to EFF-1 localization.

To visualize EFF-1 within the PLM neuron, we used a transgenic strain in which *eff-1* null mutant animals express cytoplasmic mCherry in the PLM mechanosensory neurons, as well as GFP-tagged EFF-1. We previously demonstrated that this EFF-1::GFP transgene is functional and sufficient to rescue axonal fusion defects in *eff-1* mutants^8^. Using confocal microscopy, we characterized EFF-1 localization in the cell body and proximal axon of PLM (Fig. 2a-d). In the uninjured, wild-type axon, EFF-1 formed an indiscriminate punctate pattern as we previously reported^8^ (Fig. 2b). In the cell body, it was also present as intracellular puncta, with no clear localization to the cell membrane (Fig. 2d). We chose to perform subsequent localization studies specifically in the cell body, as this wider structure allows for any mobilization of EFF-1 to the membrane to be clearly identified.

**Figure 2.**
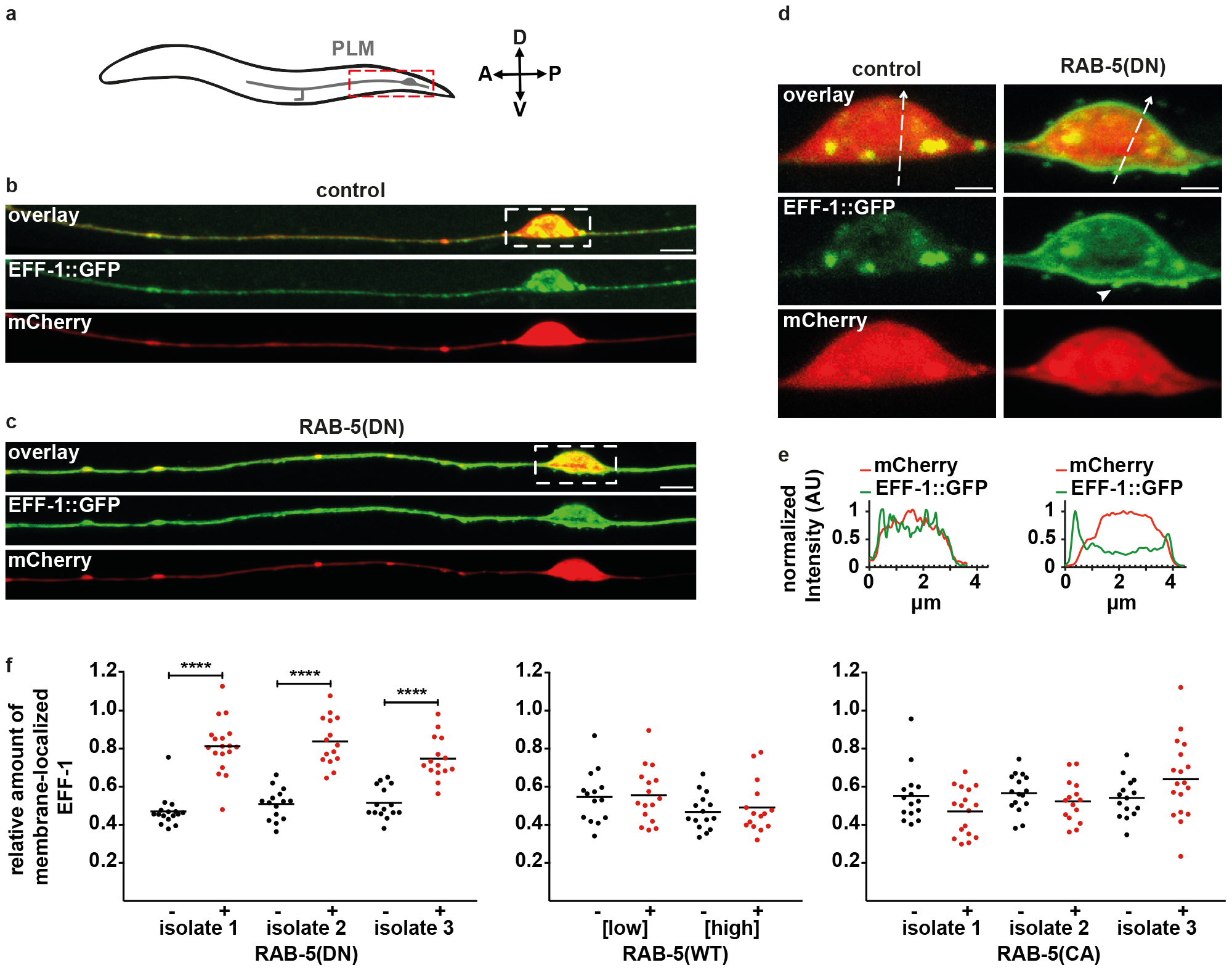
RAB-5 controls EFF-1 localization to the plasma membrane. **a** Schematic of the PLM neuron in the tail of the animal; dashed box indicates the region depicted in panels (b) and (c). **b, c** Representative maximum projection confocal images of the PLM neuron in *eff-1(ok1021)* animals expressing EFF-1::GFP and cytoplasmic mCherry in PLM. An animal expressing RAB-5(DN) (c) is compared with a sibling lacking this transgene (b). Dashed boxes indicate the regions magnified in panel (d). Scale bars 5 μm. **d** Magnification of the PLM cell bodies in panels (b) (left) and (c) (right), where EFF-1::GFP localizes to a membranous protrusion (arrowhead). Contrast settings have been adjusted to better visualize EFF-1 in the cell body rather than the axon. Scale bars 1 μm. **e** Fluorescence profiles from line scans of the overlying cell bodies confirm membranous EFF-1::GFP localization in the presence of RAB-5(DN). Results demonstrative of ≥10 animals per group. AU = arbitrary units. **f** Quantification of the relative amount of EFF-1::GFP at the membrane of the cell body in *eff-1(ok1021)* animals co-expressing EFF-1::GFP with either RAB-5(DN), RAB-5(WT) (low (5 ng/μl) and high (10 ng/μl) injection concentrations are shown) or RAB-5(CA). For each independent transgenic isolate, animals carrying a particular RAB-5 transgene (red) are compared with siblings lacking the transgene (black). Each data point represents one animal; n ≥ 15. Bars represent the mean of each group; P values from t-test: ****P<0.0001.

To test if perturbing RAB-5 activity led to EFF-1 localization at the membrane, we coexpressed RAB-5(DN). In these animals, EFF-1::GFP formed a more continuous pattern along the axon (Fig. 2c), and accumulated at the membrane of the PLM cell body (Fig. 2d), a process that was commonly associated with membranous protrusions of EFF-1 (Fig. 2d, arrowhead). Line scan profiles through these protrusions clearly demonstrated that EFF-1 was present on the membrane of the cell body, a phenomenon that we never observed in control animals (Fig. 2e). To quantify EFF-1 localization, we measured the average intensity of GFP in the membrane of the cell body, normalized to the average intensity for the whole cell body (see Methods section). This revealed a significant increase in the relative amount of EFF-1::GFP at the membrane in the presence of RAB-5(DN) (Fig. 2f). These results indicate that neuronal-specific loss of RAB-5 activity leads to EFF-1 accumulation at the membrane of the neuronal cell body.

As expected, enhancing RAB-5 activity, through either overexpression of wild-type RAB-5 or expression of RAB-5(CA), had no effect on the relative amount of EFF-1::GFP at the cell membrane (Fig. 2f). This was consistent with our axotomy results which indicated that increased RAB-5 activity did not significantly alter EFF-1 function. Taken together, these findings suggest that RAB-5 activity functions in the endocytosis of EFF-1 molecules which have been mobilized to the membrane.

### Plasma membrane accumulation of EFF-1 forms extracellular vesicles

Interestingly, we observed that animals expressing RAB-5(DN) not only had protrusions of the transmembrane EFF-1::GFP from the PLM cell body, but also generated what appeared to be extracellular EFF-1::GFP-positive vesicles. These extracellular vesicles, ranging in number from 2 - 20 per neuron, were present around the PLM cell body and proximal axon, and were reproducible in multiple independent transgenic strains (Fig. 3a). Intriguingly, previous work has shown the presence of EFF-1 vesicles in cultured medium of baby hamster kidney cells transfected with EFF-1 (ref. 27), and vesicles containing AFF-1, a second *C. elegans* fusogen, have been described both *in vitro* from mammalian cells^28^, and *in vivo* from seam cells^29^ (see Discussion).

**Figure 3.**
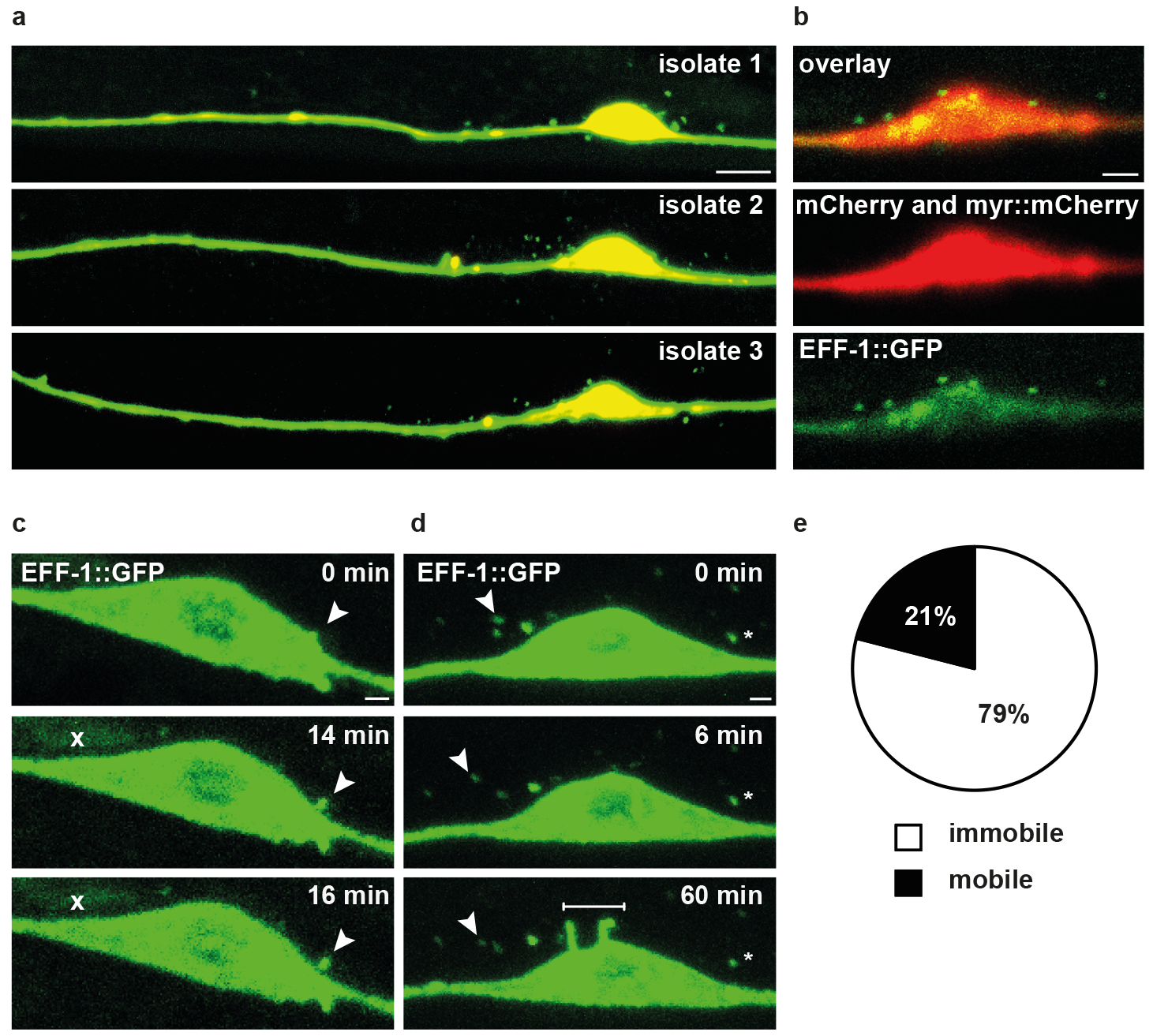
EFF-1 forms extracellular vesicles with perturbed RAB-5 activity. **a** Representative images of extracellular EFF-1::GFP vesicles in animals from three independent transgenic isolates expressing RAB-5(DN). Scale bar 5 μm. **b** Single plane confocal image of the PLM cell body in an *eff-1* mutant expressing EFF-1::GFP, RAB-5(DN), and cytoplasmic and membrane-bound mCherry (MYR::mCherry) in PLM. There is no mCherry signal present in the vesicles. Image is demonstrative of 18 animals. Scale bar 2 μm. **c** Time-lapse images demonstrating budding of an EFF-1::GFP vesicle (arrowhead). Each image represents the maximum projection of several slices to optimize visualization of the vesicle. The crosses at 14 and 16 min indicate non-specific GFP expression. Scale bar 1 μm. **d** Time-lapse images demonstrating an immobile vesicle (cross), a mobile vesicle (arrowhead) and new membranous protrusions (line). Scale bar 1 μm. **e** The proportion of mobile vs. immobile vesicles, based on 105 vesicles in 14 animals.

To determine the composition of these vesicles, we generated transgenic strains that coexpress a cytoplasmic marker (mCherry) and a membrane-bound marker (MYR::mCherry) in addition to EFF-1::GFP and RAB-5(DN). Vesicles observed in these animals presented no detectable mCherry signal (Fig. 3b), indicating that membrane-bound forms of mCherry were excluded from the vesicles, and that they contained a highly reduced volume of cytoplasm (based on the resolution of confocal imaging). This suggests that these vesicles may be selective in their composition, and contain mostly EFF-1, similar to those vesicles reported in cell culture^27^.

To characterize the dynamics of the vesicles, we undertook time-lapse imaging of the PLM cell body. Confocal imaging of the vesicles at 2 min intervals for up to 60 min revealed that they fell into two categories of mobility: ‘immobile’ vesicles, representing the majority of vesicles which exhibited no movement in this time period, and ‘mobile’ vesicles, which instead exhibited linear or disorganized, oscillatory movements around the cell body (Fig. 3d, e). This occurred at an average rate of 0.2 ± 0.05 μm/min (n = 22). We observed that these vesicles are present from the first and second larval stages. Budding of the vesicles from the cell body was a rare event, but was captured on one occasion at the fourth larval stage (Fig. 3c). EFF-1 retained at the membrane also underwent dynamic changes, and the formation of *de novo* protrusions of EFF-1 from the cell body was captured within a period of minutes (Fig. 3d).

To determine whether the vesicle dynamics were altered following neuronal injury, we performed axotomies and visualized the axon and cell body at 3 h and 6 h post-axotomy. However, we observed no clear change in vesicle number or localization at these time points (Supplementary Fig. 1). Overall, our results are consistent with a model in which these structures are likely generated via accumulation of EFF-1 at the membrane, such that EFF-1 is ‘pinched off’ in the form of a vesicle.

### RAB-5 controls EFF-1 localization to intracellular compartments

We next asked whether RAB-5 also controls EFF-1 localization to intracellular puncta. Given that EFF-1 co-localizes with RAB-5 in other cell types^14^, we hypothesized that the EFF-1::GFP puncta in PLM neurons represent EFF-1 contained in early endosomes, and are formed through RAB-5-mediated endocytosis and endosome fusion. Changes in RAB-5 activity were therefore expected to alter the morphology of these puncta.

To test this, we expressed the three different versions of RAB-5 described above and characterized the size of the EFF-1::GFP puncta present in the PLM cell body. We used the Squassh imaging tool (ImageJ)^30^ to automatically select and measure these puncta. Our results revealed no significant difference in the average size of the puncta detected in RAB-5(DN) cell bodies (Fig. 4a). However, overexpression of RAB-5(WT) caused accumulation of EFF-1 within enlarged intracellular puncta (Fig. 4b). This effect was even more pronounced when RAB-5 activity was increased by RAB-5(CA) (Fig. 4c). This result is consistent with the known role of RAB-5 in endosome fusion and the enlarged endosome phenotype generated by RAB-5 overactivity^25,26^.

**Figure 4.**
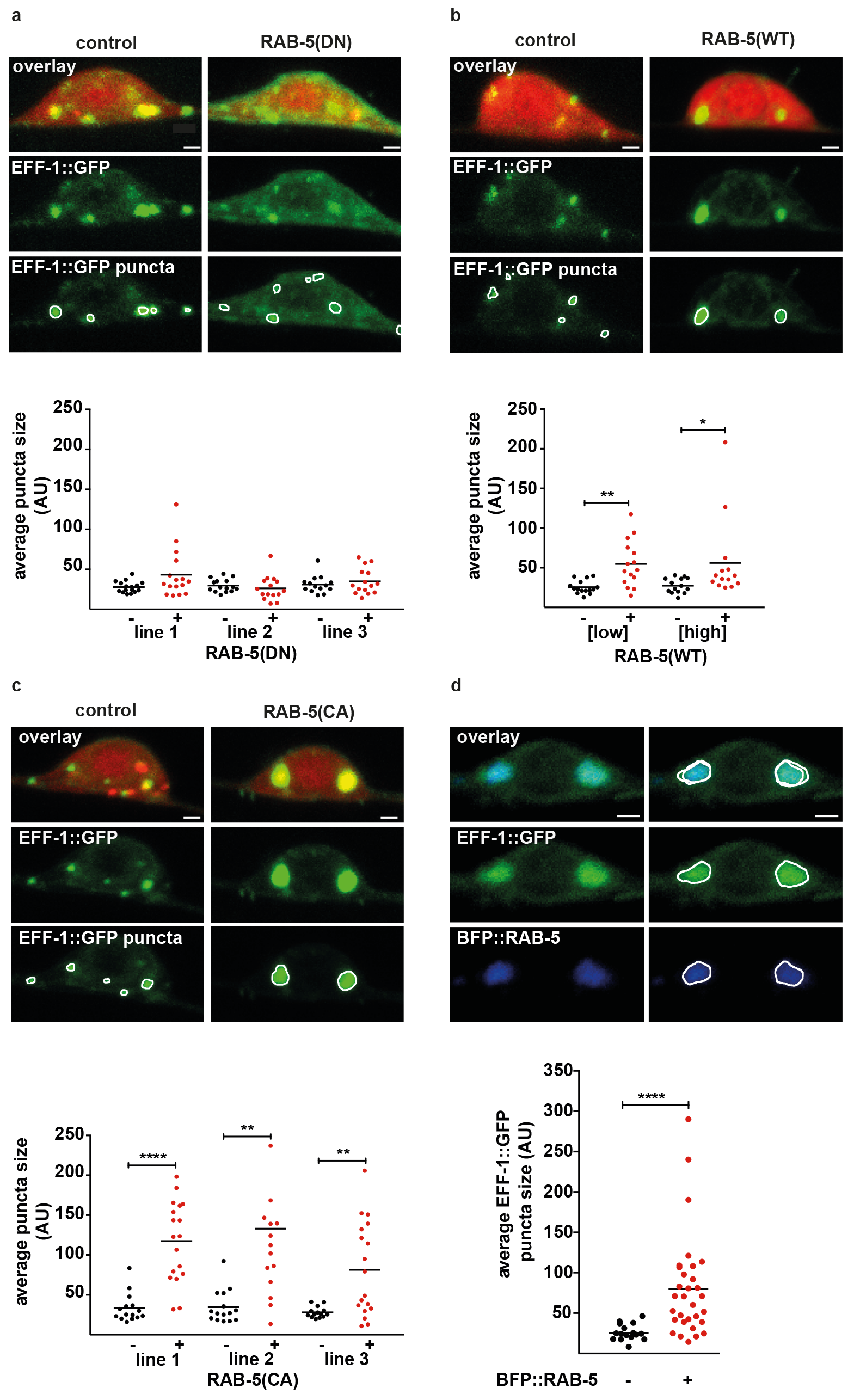
Increased RAB-5 activity causes EFF-1 accumulation in large intracellular puncta, which are RAB-5-positive compartments. **a, b, c** Representative maximum projection confocal images of the PLM cell body in *eff-1(ok1021)* animals expressing RAB-5(DN) (a), RAB-5(WT) (b) or RAB-5(CA) (c) compared with control animals lacking each respective RAB-5 transgene. Displayed for each cell body are the overlay of the red (diffusible mCherry) and green (GFP-tagged EFF-1) channels (top row), the green channel (middle row) and the result of automatic puncta selection using Squassh (bottom row). Scale bars 1 μm. Below the confocal images are graphs showing quantification of the average EFF-1::GFP puncta size in independent transgenic isolates expressing each version of RAB-5: RAB-5(DN) (a) RAB-5(WT) (b) or RAB-5(CA) (c). Animals with a given RAB-5 transgene (red) are compared with siblings lacking the transgene (black). A significant increase in average puncta size was found for three independent RAB-5(CA) isolates, and for two RAB-5(WT) isolates injected at different concentrations (5 ng/μl and 10 ng/μl). Each data point represents one cell body from one animal; n ≥ 14. Bars represent the mean of each group; P values from t-test: *P<0.05; **P<0.01, ****P<0.0001. AU = arbitrary units. **d** Representative maximum projection confocal image of the PLM cell body in an animal co-expressing EFF-1::GFP and BFP::RAB-5; the overlay and individual green and blue channels (left column) are matched with the result of automatic puncta selection using Squassh (right column), demonstrating strong co-localization between the EFF-1::GFP and BFP::RAB-5 puncta. Scale bar 1 μm. Below these confocal images is a graph showing quantification of the average EFF-1::GFP puncta size in a transgenic strain carrying EFF-1::GFP and BFP::RAB-5. A significant increase in EFF-1::GFP puncta size occurred when BFP::RAB-5 was also present (indicated by a plus sign (+)). The BFP::RAB-5 transgene was injected at the same concentration as the other RAB-5 transgenes. Each data point represents one cell body from one animal; n ≥ 17. Bars represent the mean of each group; P values from t-test: ****P<0.0001. AU = arbitrary units.

To confirm that EFF-1 was accumulating in RAB-5-positive compartments, we performed co-localization studies of EFF-1 with RAB-5 in the context of RAB-5 overexpression. We co-expressed BFP::RAB-5 with EFF-1::GFP in the mechanosensory neurons (Fig. 4d), which reproduced the enlarged EFF-1 puncta phenotype, leading to a significant increase in average EFF-1 puncta size (Fig. 4d). In these enlarged puncta, the average co-localization of EFF-1 with RAB-5 was 85% (range 67 - 98%; n = 12). Taken together, these results strongly support the notion that overactivity of RAB-5 results in EFF-1 accumulation in enlarged early endosomes.

### RAB-5 controls the amount of EFF-1 in the PLM neuron

Our results indicated that altering RAB-5 activity causes EFF-1 mis-localization to specific subcellular compartments (either the membrane with decreased RAB-5 activity, or enlarged endosomes with increased RAB-5 activity). We next asked whether this mis-localization affected the ability of the neuron to mediate recycling or degradation of the EFF-1 protein. We hypothesized that defects in such processes would result in a buildup of EFF-1::GFP in the neuron. To test this, we measured the average EFF-1::GFP intensity in the PLM axon and cell body of animals expressing either RAB-5(DN), RAB-5(CA) or RAB-5(WT). We also used an alternative approach to perturb RAB-5 activity using cell-specific RNAi^31^, whereby animals expressed *rab-5(sas)* in the mechanosensory neurons for cell-specific silencing of *rab-5*. We found that the average EFF-1::GFP intensity in both the axon and cell body was significantly greater in animals expressing either RAB-5(DN), RAB-5(CA) or *rab-5(sas)* (Supplementary Fig. 2a-d, g, h). This may represent defects in recycling and/or degradation of EFF-1 due to altered RAB-5 activity. In contrast, the overexpression of RAB-5(WT) had no significant effect on EFF-1::GFP intensity (Supplementary Fig. 2e, f). This indicates that the RAB-5(WT) molecule may be modified by endogenous regulators, and hence regulated to minimize changes in EFF-1 levels. Thus, only unregulated alterations in RAB-5 activity (generated with expression of RAB-5(DN), RAB-5(CA) or *rab-5(sas)*) disrupt EFF-1 protein levels.

### Other endocytic molecules do not control EFF-1 localization

A number of molecules function in the endocytosis of cargo from the plasma membrane. We sought to determine whether other endocytic regulators might act in conjunction with RAB-5 to regulate neuronal EFF-1. One important candidate for this role is DYN-1, the *C. elegans* orthologue of dynamin, as it has been demonstrated to act in this fashion alongside RAB-5 to regulate EFF-1 in the hypodermis^14^. We tested whether loss of DYN-1 function, induced using the temperature-sensitive allele *dyn-1(ky51)*, influenced EFF-1::GFP localization in the PLM neurons. Surprisingly, we found no significant effect of DYN-1 on EFF-1::GFP puncta size, membrane localization or average intensity in the neuron (Supplementary Fig. 3a, c, e).

We also tested EHS-1, a clathrin adaptor involved in endocytosis that localizes to the plasma membrane. However, loss-of-function *ehs-1* mutant animals showed no significant change in the same measurements of EFF-1::GFP localization (Supplementary Fig. 3b, d, f). We believe that this is consistent with studies of *ehs-1* in other systems which indicate it can act redundantly with other clathrin adaptors^32^. As such, identifying its potential role in this process requires further investigation.

From early endosomes, the endocytic pathway allows for transport of cargo to downstream compartments, potentially for membrane recycling or degradation. We therefore asked whether molecules localizing to these compartments also participated in EFF-1 regulation. A number of RAB proteins presented good candidates; these included RAB-7, which functions in endosome-to-lysosome trafficking, RAB-10, an endocytic recycling regulator that localizes to endosomes and Golgi, and RAB-11, which controls transport between recycling endosomes and the plasma membrane. We perturbed the function of these *rab* genes in the PLM neurons, either through expression of dominant negative versions, or by using loss-of-function alleles. However, our results indicated that these RAB molecules were not involved in EFF-1 localization in the PLM neurons. Neither expression of dominant negative RAB-7 or RAB-11, nor the presence of the *rab-10(dx2)* loss-of-function allele, significantly altered our measurements of EFF-1::GFP intensity or localization (Supplementary Fig. 4). Overall, these results support the specificity of our findings with RAB-5, and suggest that this molecule is a key regulator of neuronal EFF-1 and of axonal fusion.

## DISCUSSION

Our data demonstrate that perturbing RAB-5 activity in the PLM neurons has a clear functional effect on axonal repair. Specifically, the presence of RAB-5(DN) increased the capacity for EFF-1 to mediate axonal fusion, as it phenocopied EFF-1 overexpression by rescuing the *psr-1* axonal fusion defect. By visualizing EFF-1, we determined that this change in activity reflected mobilization of EFF-1 to the plasma membrane, which also occurs with reduced RAB-5 function in the *C. elegans* hypodermis^14^.

Interestingly, the presence of RAB-5(CA) did not produce the opposite phenotype, as it did not significantly reduce axonal fusion rates (Fig. 1e) or remove greater amounts of EFF-1 from the neuronal membrane (Fig. 2). It is possible that there was insufficient overactivity of RAB-5 in the strains tested to fully remove EFF-1 from the membrane, or that only a small number of EFF-1 molecules is required at the membrane for fusion, and sufficient amounts were present even in the presence of RAB-5(CA). Alternatively, it is possible that a RAB-5-independent mechanism exists for the mobilization of EFF-1 to the membrane after injury. Recruitment of EFF-1 to fusion sites in larval hypodermal cells has been shown to be mediated at least in part by the actin regulator VAB-10 (ref. 33), although other pathways may also exist. Much has been characterized about the molecular cascades that are activated in regenerating axons^34–36^ and it is plausible that some of these molecules play a currently uncharacterized role in EFF-1 recruitment.

However, overactivity of RAB-5 did lead to accumulation of EFF-1 in enlarged RAB-5-positive compartments (Fig. 4c, d). Previous studies have documented 45 - 69% colocalization of endogenous EFF-1 with RAB-5 in hypodermal cells^14^. As our model visualizes overexpression of RAB-5, our results are not directly comparable and we are unable to conclude to what extent EFF-1 co-localizes with wild-type RAB-5 levels in the neuron. However, our findings are consistent with a role for RAB-5 in determining steady-state EFF-1 localization to early endosomes.

Altering RAB-5 activity also increased the intracellular amount of EFF-1, as reflected by increases in EFF-1::GFP intensity. This occurred in both the axon and the cell body, indicating that it likely represents protein accumulation rather than an axonal transport defect. It was also observed specifically in the presence of RAB-5 molecules with locked activity states, which were associated with abnormal accumulation of EFF-1 either at the membrane or in early endosomes. There is good precedence for RAB-5 functioning in protein degradation and homeostasis. In the *C. elegans* embryo, this molecule is required for both endocytosis and degradation of the *C. elegans* caveolin CAV-1 (ref. 37). In HeLa cells, either increasing or decreasing RAB-5 activity (achieved indirectly using a regulator of RAB-5) was found to perturb endocytic trafficking and lead to cargo build-up in specific compartments^38^. We propose that the accumulation of EFF-1, either at the membrane with RAB-5(DN) or in abnormally enlarged endosomes with RAB-5(CA), occurred due to defects in the trafficking of EFF-1 from these structures. Interestingly, greater intracellular levels of EFF-1 *per se* did not guarantee an improvement in axonal fusion rates. Rather, our results indicate that EFF-1 must specifically accumulate at the membrane to improve fusion capacity.

Our findings are consistent with a model in which RAB-5 mediates the endocytosis of EFF-1 from the PLM cell membrane, similar to that described in the hypodermis^14^. In our model (Fig. 5), EFF-1 is transiently inserted into the plasma membrane following synthesis. Active RAB-5 is responsible for subsequent transport of this EFF-1 into early endosomes, where EFF-1 largely resides in the steady state. Decreasing RAB-5 activity allows EFF-1 accumulation at the membrane, whereas overactivity of RAB-5 leads to accumulation in early endosomes. We also propose that, in neurons, RAB-5-mediated endocytosis occurs upstream of pathways for EFF-1 recycling/degradation. Because altering RAB-5 activity sequesters EFF-1 in specific compartments, it prevents downstream trafficking and creates a build-up of EFF-1 in the neuron.

**Figure 5.**
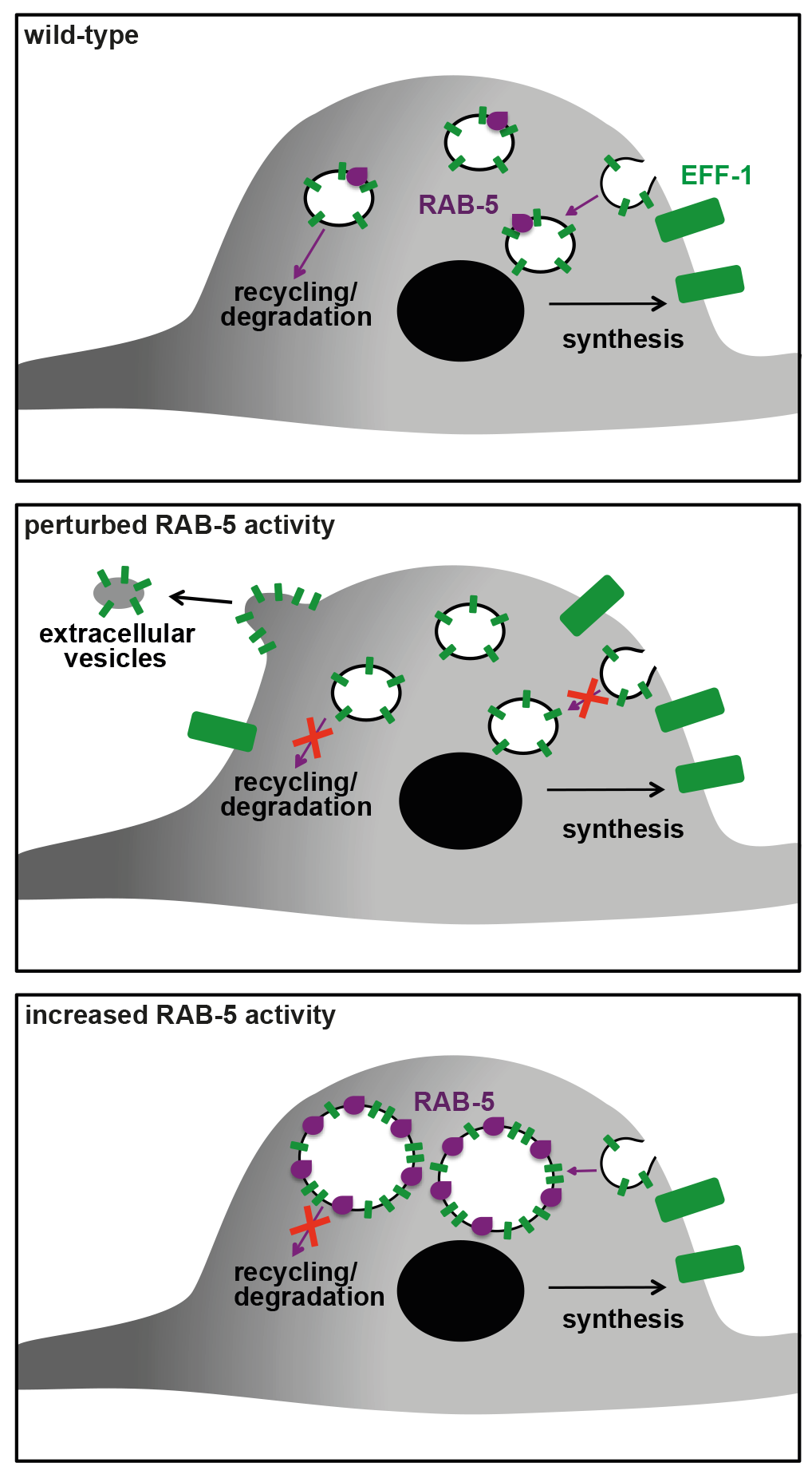
Model of RAB-5 regulation of EFF-1 in PLM. In the wild-type scenario, EFF-1 (green) is transiently inserted into the membrane following synthesis. Active RAB-5 (purple) functions in removal of this EFF-1 into early endosomes. This in turn is required for EFF-1 trafficking into downstream compartments for recycling/degradation. When RAB-5 function is perturbed, EFF-1 is no longer removed from the membrane, and subsequent accumulation at the membrane results in membranous protrusions and budding of extracellular vesicles. There is an additional failure of trafficking into recycling/degradation pathways. With increased RAB-5 activity, EFF-1 instead accumulates in enlarged early endosomes. This endosomal enlargement also leads to a defect in downstream trafficking of EFF-1 for recycling/degradation.

A particularly fascinating aspect of this study was the generation of extracellular EFF-1::GFP-positive vesicles when RAB-5 activity was perturbed. Extracellular vesicles were observed incidentally in the earliest studies of EFF-1-mediated cell-cell fusion using electron microscopy^39^. Vesicles that specifically contain fusogens have now been documented *in vitro*^27,28^, and more recently *in vivo* in *C. elegans*^29^, but their exact characteristics and functionality are yet to be elucidated. The absence of both membrane and cytoplasmic markers in the EFF-1::GFP vesicles suggests that these vesicles may exclude some membrane proteins, such as fluorophores, or contain insufficient amounts for visualization with confocal microscopy. However, it appears unlikely that they contain purely EFF-1::GFP, as fusogen-containing vesicles *in vitro* have been shown to contain other proteins^27^, as well as membrane proteins in some cases^28^. We therefore postulate that these vesicles may be selective in their uptake of membrane proteins, and that their content could differ from the original composition of the PLM plasma membrane. Given that extracellular vesicles are known to deliver cargo in diverse systems^40^, fusogen-containing vesicles represent an attractive vehicle for imparting fusion competence to surrounding tissues, and potentially to mammalian neurons, as previously postulated^29^. The evidence that these vesicles may be selective, and largely contain fusogen, suggests that they could be very efficient in delivering fusogenicity, although whether the EFF-1::GFP vesicles have fusogenic activity is currently unclear.

Another matter of speculation is the mechanism through which the EFF-1::GFP vesicles are generated. It remains to be determined whether they are passively extruded, or are instead actively secreted. Given that they occur in the presence of increased intracellular and membranous EFF-1, it is plausible that they are created through excessive build-up of EFF-1 at the membrane. With its known function in membrane sculpting^41^, EFF-1 could potentially ‘pinch off’ a section of membrane. Consistent with this, the vesicles localize in the vicinity of the PLM cell body, and it may be that the higher volume-to-surface-area ratio in this part of the neuron allows for sufficient build-up of EFF-1::GFP behind the membrane. In support of this, most protrusions, and the observed event of vesicle budding, originated from the cell body. However, if there is instead molecular machinery for active secretion of these vesicles, one strong candidate is the ABC transporter CED-7, which is known to generate extracellular vesicles containing phosphatidylserine during apoptotic cell clearance^42^. We have shown that CED-7 functions in regeneration of the PLM neuron^8^ as well as its degeneration^43^, possibly through vesicle generation. It is therefore possible that CED-7 activity in the neuron is required for the secretion of EFF-1 vesicles.

RAB proteins have well-established roles in intracellular trafficking, and we tested a suite of molecules other than RAB-5 that could potentially regulate such transport of EFF-1 in PLM. As reported for the hypodermis^14^, RAB-7, RAB-10 and RAB-11 did not influence EFF-1 localization. Similarly, we did not find a role for DYN-1, although this molecule has been demonstrated to negatively regulate EFF-1 and cell-cell fusion in the hypodermis^14^. However, the literature on dynamin suggests that it can have varying roles in fusion events in different systems, related to its multiple functions in endocytosis as well as actin cytoskeletal rearrangements. In some mammalian cell-cell fusion, including in osteoclasts and myoblasts, dynamin activity appears to instead promote fusion^44^. Our result is in keeping with an alternative role for DYN-1 in the neuron, potentially in endocytic regulation of molecules other than EFF-1.

In summary, our study identifies RAB-5 as a key regulator of EFF-1 in the nervous system. By controlling the levels of EFF-1 in endosomal compartments, RAB-5 modulates the amount of this protein available on the membrane, thereby regulating its fusogenic capacity. Thus, manipulating the activity of RAB-5 activity provides a means to promote highly efficient neuronal repair through axonal fusion.

## METHODS

### Strains and genetics

Standard techniques were used for *C. elegans* strain maintenance and genetic manipulations^45^. All experiments were performed at 22 °C (room temperature) on L4 animals unless otherwise specified. The following mutations were used: *rab-10(dx2) I, eff-1(ok1021) II*, *ehs-1(ok146) II, psr-1(ok714) IV, dyn-1(ky51) X*. The integrated transgenic strain QH3135 *[zdIs5(Pmec-4::GFP) I]* was used as a background strain for performing axotomies. The transgenic strain QH4748 *[eff-1(ok1021) II; vdEx662[Pmec-4::eff-1::gfp; Pmec-4::mCherry; Podr-1::DsRed]]*^8^ was used as a background strain for all experiments involving EFF-1 confocal imaging. To generate extrachromosomal arrays, microinjections were performed into the germline using standard methods^46^. The following transgenes were generated (concentrations used for the microinjection mix are indicated in brackets; all injection mixes had a total concentration made up to 100 ng/μl using empty pSM plasmid): *vdEx1192/vdEx1193/vdEx1197[Pmec-3::rab-5(S33N) (*5 ng/μl); *Podr-1::gfp* (60 ng/μl)*], vdEx1450/vdEx1451/vdEx1452[Pmec-3::rab-5(Q78L)* (5 ng/μl); *Podr-1::gfp* (60 ng/μl)*], vdEx1194 [Pmec-3::rab-5(WT)* (5 ng/μl)*; Podr-1::gfp* (60 ng/μl)], *vdEx1237[Pmec-3::rab-5(WT)* (10 ng/μl); *Podr-1::gfp* (60 ng/μl)], *vdEx1051/vdEx1055/vdEx1084[Pmec-3s::rab-5(s)* (5 ng/μl); *Pmec-3::rab-5(as) (*5 ng/μl); *Podr-1::gfp* (60 ng/μl)], *vdEx1301/vdEx1302/vdEx1303[Pmec-3::rab-7(T23N)* (5 ng/μl); *Podr-1::gfp* (60 ng/μl)], *vdEx1375/vdEx1390/vdEx1443[Pmec-3::rab-11(S25N)* (5 ng/μl); *Podr-1::gfp* (60 ng/μl)], *vdEx1389[Pmec-3::bfp::tev-s::rab-5* (5 ng/μl)*; Pmyo-2::mCherry* (2.5 ng/μl)], *vdEx1566[Pmec-4::myr::mCherry* (5 ng/μl)*; Pmyo-2::mCherry* (2.5 ng/μl)], *vdEx1576 [Pmec-4::myr::mCherry* (15 ng/μl)*; Pmyo-2::mCherry* (2.5 ng/μl)]. It should be noted that a TEV-S signal exists between the BFP and RAB-5 sequences in *vdEx1566*; this signal allows for protein cleavage with addition of TEV protease and is therefore highly unlikely to have affected the outcome of the current study.

### Molecular biology

Standard molecular biology techniques were used^47^. To generate the plasmid *Pmec-3::rab-5(WT)*, the *rab-5* gene was amplified from the pCL206 plasmid using 5’-gcTCTAGAatggccgcccgaaacgcagg-3’ and 5’-gggaCCCGGGttatttacagcatgaaccc-3’. These primers introduced Xba I and Xma I restriction sites respectively, which were used to clone the amplicon into the plasmid L1026 (*Pmec-3*, from Fire Vector Kit 1995). The QuikChange II Site-Directed Mutagenesis Kit (Agilent Technologies) was then used to generate *rab-5(Q78L) (CA)* and *rab-5(S33N (DN)* variants of this plasmid, using the primers 5’-aaatctgggatactgcaggaaaagaaagatatcattattgg-3’, 5’- ccaatgaatgatatctttcttttcctgcagtatcccgattt-3’ and 5’ -ctatcatttcaggcaaaaactctctcgtattgcgattc-3’, 5’-gaatcgcaatacgagagagtttttgcctgaaatgatag-3’ respectively.

To generate *Pmec-3s::rab-5(sas)*, a sense-antisense PCR fusion technique was used^31^. *rab-5* was amplified from genomic *C. elegans* DNA using standard primers (5’-cgtgccttcaatctttttcg-3’ and 5’-acaatgacgacgatcacaggc-3’). *Pmec-3s* was amplified from the L3784 plasmid (*Pmec-3::gfp*, from Fire Vector Kit 1997) with a standard forward primer (5’-aggtacccggagtagttggc-3’) and two different reverse primers with sequences complementary to the extremities of *rab-5* at the 5’ ends (5’-atgttgcatttttctttccagaatctataacttgatagcgata-3’ and 5’-cttcccaactaccatgtacaaaatctataacttgatagcgata-3’). The *rab-5* and *Pmec-3s* reactions products were then fused using nested primers 5’-ggcagtaatgaagacgtccat-3’ and 5’-gaagggttgatggtacatgaaa-3’ or 5’-ttctggaaagaaaaatgcaacat’-3’.

*Pmec-3::rab-7(T23N) (DN)* and *Pmec-3::rab-11(S25N) (DN)* were generated using the QuikChange II Site-Directed Mutagenesis Kit on *Pmec-3::rab-7* and *Pmec-3::rab-11* plasmids respectively, with primers 5’-cgggcgttggaaagaattctttgatgaatcaatatg-3’, 5’-catattgattcatcaaagaattctttccaacgcccg-3’ and 5’-gagactcaggcgtcggaaagaataatctcctgtctcgtttcac-3’, 5’-gtgaaacgagacaggagattattctttccgacgcctgagtctc-3’ respectively. The construction of *Pmec-3::rab-7* and *Pmec-3::rab-11* involved amplification of each gene from *C. elegans* genomic DNA with primers to introduce Xba I and Xma I restriction sites, followed by cloning into the L1026 plasmid downstream of *Pmec-3*.

To generate *Pmec-3::bfp::tev-s::rab-5*, the *bfp::tev-s::rab-5* insert was amplified from *pOG172*, a kind gift from Prof. Guangshuo Ou (Tsinghua University, Beijing). Primers were used to introduce BamH I and Msc I restriction sites for cloning into L1026 downstream of *Pmec-3*. To generate the *Pmec-4::myr::mCherry* plasmid, *myr::mCherry* from the *PNV::myr::mCherry* plasmid was cloned into *Pmec-4::GFP* using Msc I and EcoR I restriction enzymes which remove the GFP sequence.

### Laser axotomy

We performed UV laser axotomy of PLM in animals at the L4 larval stage as previously described^7,8^. Animals were anaesthetized using 0.05% tetramizole hydrochloride on 4% agarose pads. The axotomy was performed approximately 50 μm from the cell body using a MicroPoint Laser System Basic Unit attached to a Zeiss Axio Imager A1. At 48 h post-axotomy, animals were analyzed on a Zeiss Axio Imager Z1 equipped with a Photometrics Cool Snap HQ2 camera with MetaMorph software for the presence of reconnection and fusion. If it was unclear whether a distal fragment had been maintained, the animal was scored again at 72 h post-axotomy.

### Confocal microscopy

Localization studies of EFF-1 and RAB-5 were performed on a LSM 710 META confocal microscope, equipped with a GaAsP detector and Zen 2012 software. L4 animals were mounted on 3% agarose pads in 25 mM sodium azide. Separate *Z*-stacks were performed of the PLM cell body and proximal axon. For imaging of EFF-1::GFP and cytoplasmic mCherry, green fluorescence was visualized with a 488 nm laser (5% power for the axon, 2% power for the cell body; gain of 600 and 4x averaging for both) and red fluorescence was visualized with a 543 nm laser (1% power for the axon, 0.2% power for the cell body; gain of 500 and 4x averaging for both). For imaging of BFP::TEV-S::RAB-5, blue fluorescence was visualized with a 405 nm laser (0.5% power, gain of 500, 4x averaging). To image MYR::mCherry, red fluorescence was again visualized with a 543 nm laser (up to 10% power, gain of 500, 4x averaging).

To image EFF-1::GFP post-axotomy, animals were mounted for axotomies in tetramizole as described above. They were then recovered in drops of M9 buffer onto seeded NGM plates for either 3 h or 6 h, after which they were mounted in sodium azide for confocal imaging.

### Vesicle time lapse imaging

For characterization of EFF-1::GFP vesicle dynamics, *z*-stacks of the PLM cell body were acquired at 30 s or 2 min intervals for a total of 6 - 60 min depending on the dynamics observed. Vesicles were classified as immobile or mobile based on the presence of any movement in this timeframe. Mobile vesicles (22 out of 105) were measured for movement in the XY axis per frame by drawing a linear region of interest (ROI) from the center of the vesicle to the center at its subsequent location; the total distance was averaged over the time imaged to be expressed in μm/min.

### *dyn-1* heat-shock

To test the temperature-sensitive allele *dyn-1(ky51)*, heat-shocks were performed for either 30 min or 2 hr. L4 animals were placed on NGM plates in a 25 °C incubator. Mutant and control animals were heat-shocked concurrently on separate plates and mounted on the same slide for subsequent confocal imaging. *dyn-1(ky51)* animals raised at 25 °C did not lay viable eggs, whereas those laid at 15 °C and then transferred to 25 °C developed to the L4 stage. However, the animals raised at 25 °C demonstrated non-specific increases in EFF-1::GFP intensity in both mutant and control groups (data not shown). Heat-shocks used in this assay were therefore limited to a maximum of 2 h.

### Confocal image analysis

Image analysis was performed using Fiji for Mac OS X (ImageJ). To score for the presence of EFF-1::GFP on the membrane of the cell body, fluorescence profiles of line scans were obtained using the ‘Plot Profile’ tool in ImageJ. EFF-1::GFP was scored as localizing to the membrane if the peak of green intensity (EFF-1::GFP) and the peak of red intensity (cytoplasmic mCherry) did not overlap.

For EFF-1::GFP intensity calculations, average projections of *z*-stacks were analysed. For the axon, a line scan was performed along the axon for the initial ~50 μm anterior to the cell body, and the mean intensity in the green and red channels was measured. Background fluorescence was calculated using the same line scan moved to three different positions around the axon in the image; the mean background fluorescence from these three readings was subtracted from the average intensity value of the axon. The ratio of the GFP intensity to the mCherry intensity was then calculated to control for differences in transgene expression between animals. For intensity calculations of the cell body, a region of interest was drawn following the boundary of the cell body, and calculations were performed as for the axon.

For measurements of EFF-1::GFP border localization, average projections of *z*-stacks were analysed. A region of interest was drawn along the border of the PLM cell body. The average GFP intensity was first measured along this line (border intensity), after which the average GFP intensity was measured within the full area (cell body intensity). The final measurement was expressed as a ratio of the average border intensity to cell body intensity. Background subtraction was performed as described for intensity measurements.

To quantify EFF-1::GFP puncta size and number, maximum projection confocal images of *z*-stacks were analysed. Puncta were identified automatically using the ImageJ plugin Squassh^30^. The following settings were used: background removal, rolling ball window size 10, regularization 0.1, minimum object intensity 0.3, subpixel segmentation, automatic local intensity estimation, Poisson noise model, and Gaussian psf approximation as for confocal microscopy. The circularity of the resulting objects was calculated using the ‘Analyse Particles’ function on ImageJ. The following exclusion criteria were applied to all objects: circularity <0.7, and object located in the proximal axon or outside the neuron. If none of the puncta selected in a cell body met these criteria, the cell body was discarded from the analysis.

For co-localization studies of EFF-1 and RAB-5, only enlarged puncta were analysed; these were defined as puncta >125 units, as puncta of this size were specific to RAB-5 overactivity and never observed in non-transgenic controls. Squassh analysis was applied to both the green and blue channels, providing co-localization values for the selected objects. Identical settings (described above) were used for the two channels.

### Statistical testing

Statistical analysis was performed using GraphPad Prism and Microsoft Excel. Error of proportions was used to assess variation across a single population. Two-way comparison was performed using either the t-test or t-test with Welch correction (the latter was performed if the standard deviations of two compared groups were significantly different).

## ACKNOWLEDGMENTS

We thank Luke Hammond and Rumelo Amor for help with microscopy, as well as Rowan Tweedale and members of the Hilliard lab for helpful discussion and comments. We also thank Guangshuo Ou for the pOG172 plasmid, Nick Valmas for the *PNVMYR::mCherry* plasmid, and Elia Di Schiavi for advice regarding cell-specific RNAi. This research was supported by a University of Queensland Research Scholarship to C.L and a HFSP Fellowship LT000762/2012 to R.G.S. Some strains were provided by the *Caenorhabditis* Genetic Center (CGC), which is funded by NIH Office of Research Infrastructure Programs (P40 OD010440). This work was supported by NHMRC project grants (1068871 and 1129367), an ARC discovery project (160104359) and an NHMRC Senior Research Fellowship (1111042) to M.A.H.; NHMRC project grants (1101974 and 1099690) to B.N. The LSM 710 META confocal microscope used in this study was supported by an ARC Grant (LIEF LE130100078).

## AUTHOR CONTRIBUTIONS

C.L. designed and performed experiments and wrote the paper; B.N. provided reagents and interpreted experiments; R.G.S. and M.A.H. designed and interpreted experiments and wrote the paper.

---

**Supplementary Figure 1.**
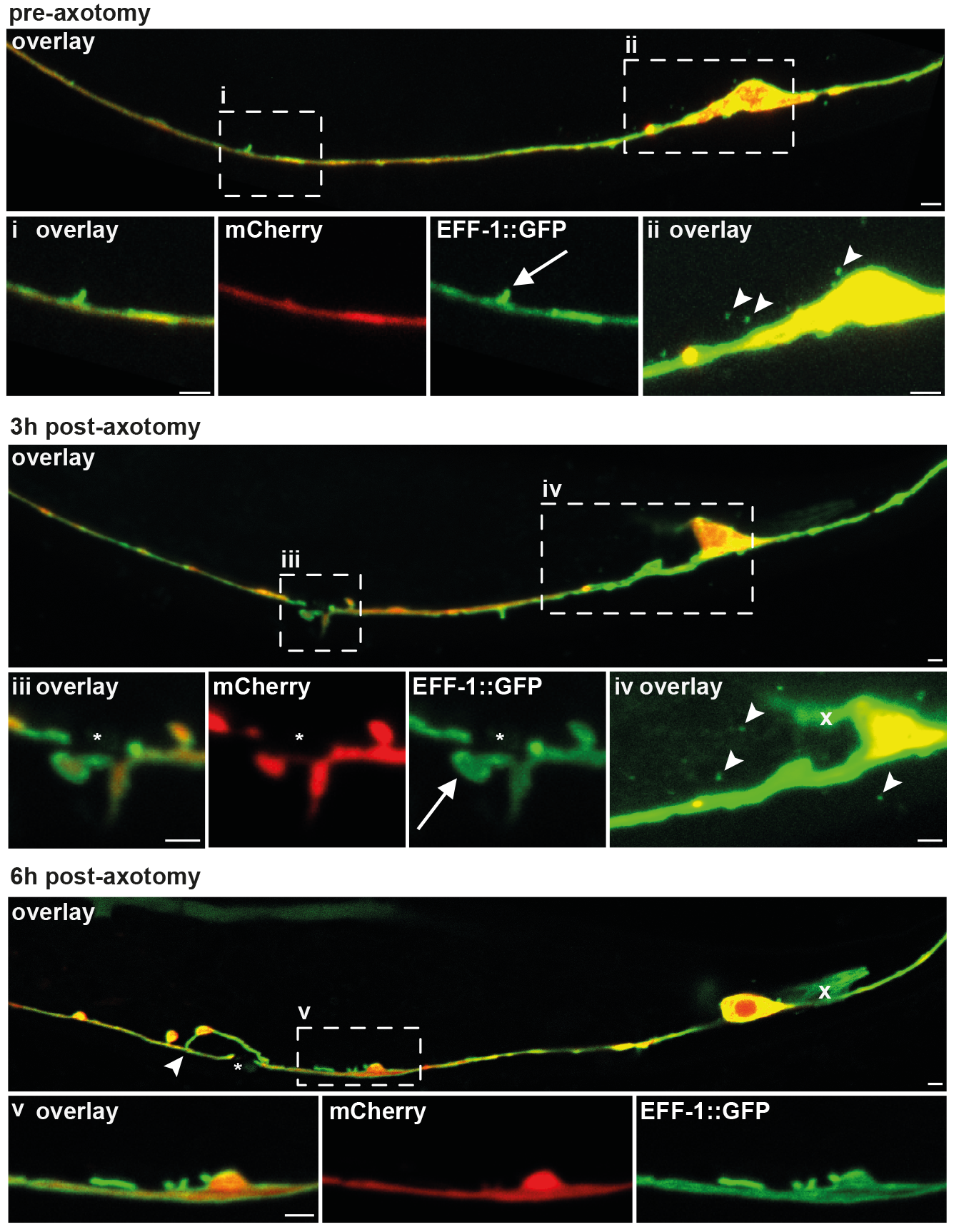
EFF-1 localization pre- and post-axotomy with perturbed RAB-5 activity. Representative confocal images of the PLM cell body and axon in *eff-1(ok1021)* animals expressing EFF-1::GFP, mCherry and RAB-5(DN). Three different animals are shown. Pre-axotomy, EFF-1 in the axon forms a continuous localization pattern, and can be visualized in areas on the membrane (box i, arrow marks a membranous protrusion). Vesicles are visualized around the cell body (box ii, arrowheads mark individual vesicles). At 3 h post-axotomy, the regenerating axon has formed a growth cone lined with EFF-1::GFP (box iii, asterisk indicates axotomy site; arrow indicates growth cone). EFF-1::GFP vesicles are present in the same number and location as prior to injury (box iv, arrowheads mark individual vesicles, cross indicates a membranous protrusion of EFF-1 that is out of focus). At 6 h post-axotomy, EFF-1::GFP can be seen forming membranous protrusions (box v) on a regenerating axon that has undergone fusion. Asterisk indicates the axotomy site; arrowhead indicates the site of fusion. A cell behind PLM non-specifically expresses GFP (indicated by cross). Images are representative of >20 animals pre-axotomy, 14 animals at 3 h post-axotomy, and 9 animals at 6 h post-axotomy. Scale bars 2 μm.

**Supplementary Figure 2.**
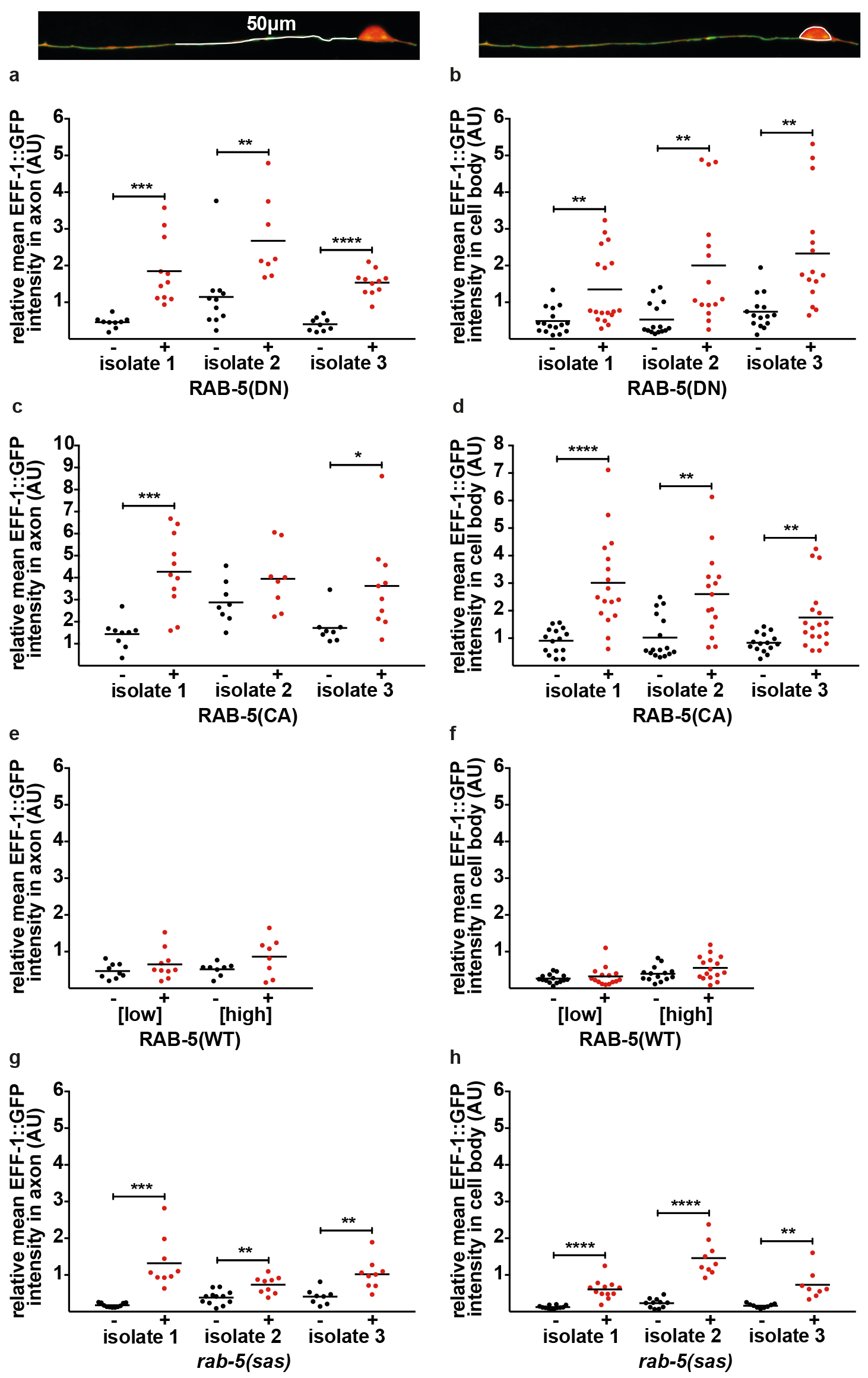
Altered RAB-5 activity increases the EFF-1::GFP intensity in PLM. Quantification of the relative mean EFF-1::GFP intensity in the axon (left column) and cell body (right column) of *eff-1(ok1021)* animals co-expressing EFF-1::GFP with different RAB-5 transgenes. ‘Isolate’ refers to an independent transgenic strain. Each mean GFP measurement is expressed relative to the mean mCherry intensity in the same region (see Methods). This intensity was significantly increased with expression of RAB-5(DN) (**a, b**), RAB-5(CA) (**c, d**), or *rab-5(sas)* (**g, h**). No significant difference was found with expression of RAB-5(WT) (**e, f**). Each point represents the mean for a single axon or cell body; n ≥ 8. Bars represent the mean of each group; P values from t-test: *P<0.05; **P<0.01; ***P<0.001; ****P<0.0001. AU = arbitrary units.

**Supplementary Figure 3.**
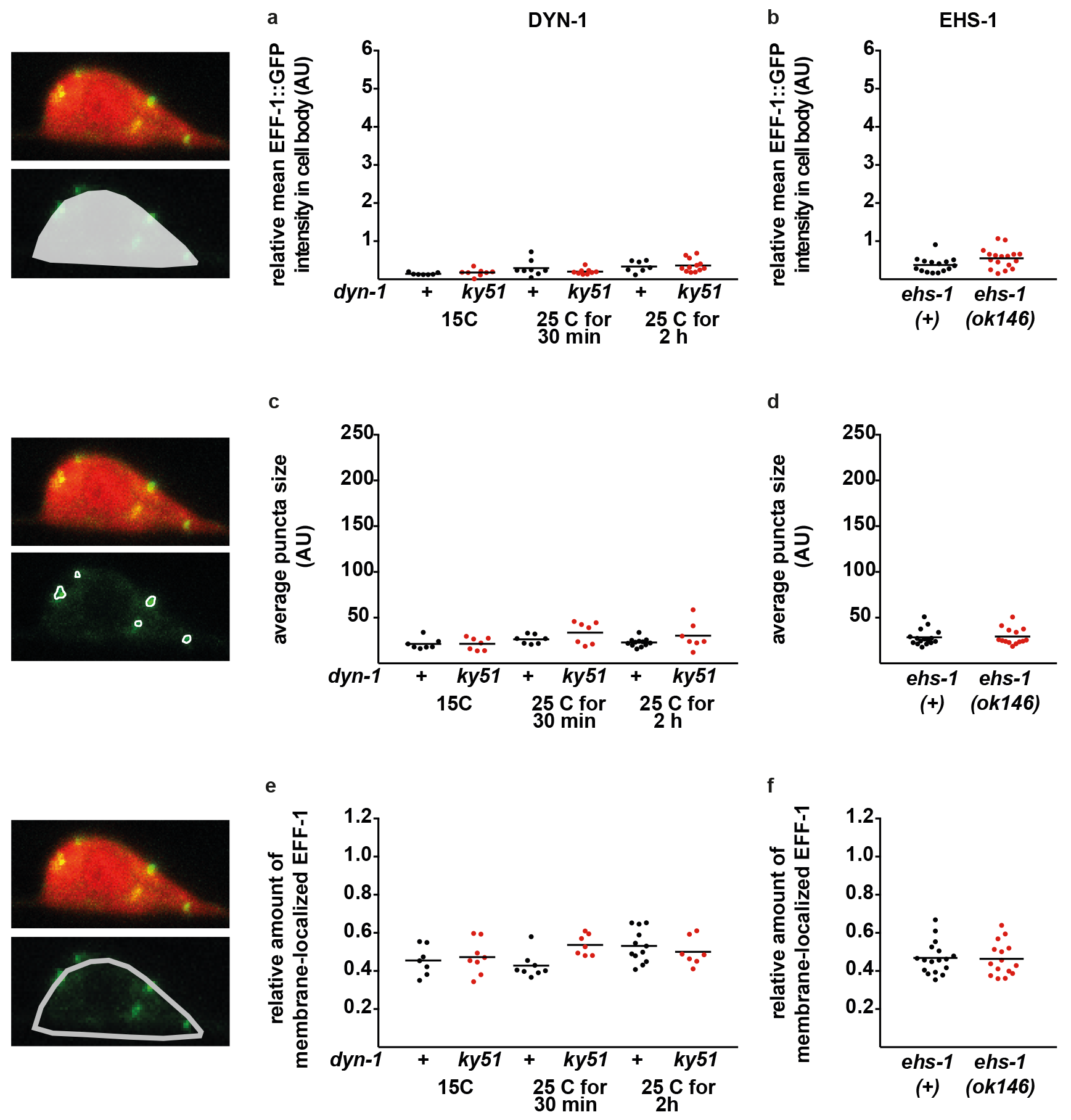
The endocytic molecules DYN-1 and EHS-1 have no effect on EFF-1 localization in PLM. Analyses of EFF-1::GFP localization in the presence of the temperature-sensitive allele *dyn-1(ky51)* (**a, c, e**) or the loss-of-function allele *ehs-1(ok146)* (**b, d, f**). No significant difference was found in EFF-1::GFP intensity (top row), the average EFF-1::GFP puncta size (middle row) or the amount of EFF-1 at the membrane (bottom row). To test *dyn-1(ky51)*, animals maintained at the permissive temperature (15 °C) are compared with those heat-shocked at the restrictive temperature (25 °C) for 30 min or 2 h. Each point represents the mean for a single cell body; n ≥ 7. Bars represent the mean; P values from t-test. AU = arbitrary units.

**Supplementary Figure 4.**
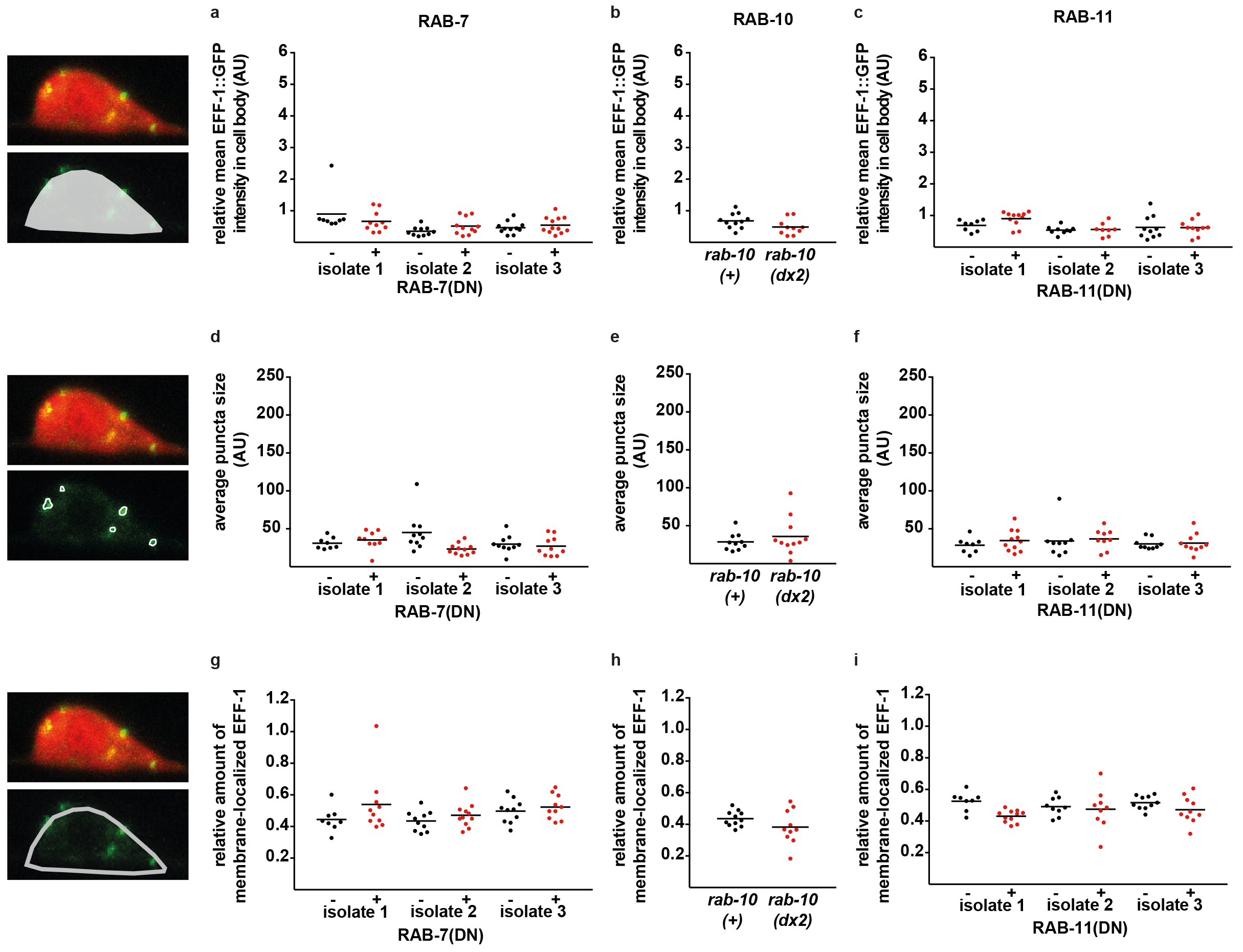
Other RAB molecules have no effect on EFF-1 localization in PLM. Analyses of EFF-1::GFP localization in independent transgenic isolates co-expressing RAB-7(DN) (**a, d, g**), RAB-11(DN) (**c, f, i**), or *rab-10(dx2)* (**b, e, h**). For each isolate tested, no significant difference was found in the relative mean EFF-1::GFP intensity (top row), average EFF-1::GFP puncta size (middle row) or amount of EFF-1 at the membrane (bottom row). Animals with the altered *rab* gene of interest (red) are compared with siblings wild-type for that *rab* gene (black). Each point represents the mean for a single cell body; n ≥ 8; bars represent the mean of each group; P values from t-test. AU = arbitrary units.

